# Phenotype-based single-cell transcriptomics reveal compensatory pathways involved in Golgi organization and associated transport

**DOI:** 10.1101/2022.12.02.518815

**Authors:** Sanjana Singh, Joanna Zukowska, Aliaksandr Halavatyi, Jonathan J. M. Landry, Rainer Pepperkok

## Abstract

The Golgi is a dynamic organelle with a unique morphology that has implications on its function. How the structural integrity of the Golgi is maintained despite its dynamic nature has been a long-standing question. Several siRNA-based screens have addressed this question and have identified a number of key players required for Golgi integrity. Interestingly, they also reported heterogeneity of phenotypic responses with regards to Golgi morphology. Although never systematically investigated, this variability has generally been attributed to poor transfection efficiency or cell cycle specific responses. Here we show that this heterogeneity is the result of differential response to the siRNA knockdown in different Golgi phenotypes, independent of transfection efficiency or cell cycle phases. To characterize the observed Golgi phenotype-specific responses at the molecular level we have developed an automated assay which enables microscopy-based phenotype classification followed by phenotype-specific single-cell transcriptome analysis. Application of this novel approach to the siRNA mediated knockdown of USO1, a key trafficking protein at the ER to Golgi boundary, surprisingly suggests a key involvement of the late endosomal/endocytic pathways in the regulation of Golgi organization. Our pipeline is the first of its kind developed to study Golgi organization, but can be applied to any biological problem that stands to gain from correlating morphology with single-cell readouts. Moreover, its automated and modular nature allows for uncomplicated scaling up, both in throughput and in complexity, helping the user achieve a systems level understanding of cellular processes.

## Introduction

Compartmentalized cells rely on the secretory and endocytic pathways to transport biomolecular cargo within and across cells^1,2^. The Golgi Complex is an indispensable part of this trafficking machinery and is a fundamental feature of eukaryotic cells^3^. Traditionally known for its functions of glycosylation, sorting and packaging of cargo^4–6^, work in recent years has demonstrated various new roles attributed to the Golgi^7,8^. It is now evident that the Golgi is involved in regulation of an ever-increasing list of cellular pathways; including but not limited to cytoskeletal remodeling^9,10^, mitosis^11,12^, apoptosis^13^, stress response^14–16^ and immune regulation^17,18^. It is likely that the forthcoming years will bring to light further functions of the Golgi as a hub for the convergence and regulation of multiple cellular pathways.

The Golgi complex presents a unique structure in the cell, comprising of flattened membrane discs (cisternae) stacked on top of each other^19^. The resulting stack is polarized, receiving newly synthesized material from the ER via the *cis* face and processed cargo leaving via the *trans* face to the TGN^20^. Additionally, in higher vertebrates, individual Golgi stacks are laterally linked to form a ribbon structure, usually stationed near the nuclear periphery^8,21^. Remarkably, this highly organized structure of the Golgi is also extremely dynamic, suggesting the presence of a large regulatory network capable of rapidly modulating Golgi structure^22^. The organelle can undergo rapid disassembly into various altered patterns, such as fragmentation, dispersion, and compaction. These altered forms can represent physiological events, such as mitosis, where the Golgi breaks down on the onset of metaphase and re-organizes itself in the two daughter cells^11,12,23^. However, altered Golgi structure could be also an indication of cellular abnormalities - such as DNA damage^24^, onset of cell death^13^, and infection^7^. Moreover, such defects have also been correlated with several diseases, including neurodegenerative diseases ^25,26^ and certain forms of cancer^27–29^. It is difficult to ascertain whether the change in Golgi structure in such cases is a cause or consequence of disease, and the link between the structural abnormalities and disease outcome is not clear^30,31^.

Golgi organization and maintenance is mediated by many contributing factors, i.e., scaffold proteins like the Golgins^32,33^ and GRASPS^34,35^; the cytoskeletal network, trafficking proteins^7,36^ as well as cargo flux through the organelle itself ^37,38^. Although much of the machinery that is involved in these processes is known, it is extremely difficult to study the structure of the Golgi in isolation of its function^30^. Several strategies have been employed to probe the structure of the Golgi morphology and ascertain the functional consequences. A popular technique is the use of drugs that disrupt Golgi structure, such as Brefeldin A – which blocks retrograde traffic^39–41^, or Nocodazole, a microtubule depolymerizing agent^40,42,43^. While such experiments have provided some key mechanistic insights, they suffer from having non-specific effects on other pathways and are quite removed from a physiological setting. Another major approach has been to knockdown key proteins and assay their importance for structural integrity of the Golgi complex. siRNA knockdowns and CRISPR based knockouts have been performed either for single proteins^44–46^ or as large/genome-wide screens^47–51^ to search for potential proteins involved in regulation of Golgi structure. While such studies have been instrumental in identifying individual components, it is difficult to judge if the effects of a candidate gene/protein thus discovered are direct or indirect.

Light & electron microscopy^52–56^, proteomics^57–59^ and mechanistic models^37,46,60–62^ have all contributed in crucial ways to the current knowledge base of Golgi architecture. One common drawback, however, of all these approaches is that they all provide single readouts (Golgi phenotype, interaction pairs, single gene expression). Maintenance of Golgi structure is likely achieved and regulated by a large, diverse system of interacting proteins, making singular readouts inefficient at unravelling entire pathways that orchestrate this.

Bulk omics (proteomics/transcriptomics) approaches are powerful tools to identify new factors and interactions, both metabolic and regulatory. These methods capture the average state of the population and assume that the real biological mechanism is represented by the average. However, it is increasingly clear that even well-established cell lines are heterogenous in nature^63^. Phenotypic heterogeneity in response to a perturbation may be of functional significance that is lost in an average measurement. Exploiting the intrinsic heterogeneity of cellular responses to perturbation is an excellent way to study the effects of a single perturbation with as few variables as possible influencing the experimental readout. To study the molecular regulation of Golgi organization in an unbiased manner, we decided to set up a method for single-cell transcriptomics of distinct Golgi phenotypes upon a treatment. The performed analysis highlighted novel insights into the functional consequences of the phenotypic differences at the network scale. Moreover, the developed pipeline can be easily adapted to correlating morphological phenotypes seen under the microscope other than the Golgi phenotype studied here with single-cell transcriptome data.

## Results

### Differences in Golgi morphology are not explained by transfection efficiency

siRNA based knockdown of certain proteins cause alterations in Golgi structure. These structural changes are not uniform across cells population, resulting a phenotypic heterogeneity observed by us and others^49,64^. This effect is usually attributed to inefficient transfection of siRNA, but this claim has never been systematically investigated before. Intrigued by this heterogeneity, we performed siRNA mediated depletion of 20 Golgi-localized proteins (Supplementary Table 1) in HeLa cells expressing a GFP-tagged version of the Golgi enzyme N-acetyl Galactosyltransferase, referred to henceforth as HeLa GalNaC-GFP. After siRNA treatment, cells were fixed and stained with the antibody against the targeted protein. The images were first analyzed for the effect on Golgi morphology, and 8 of the 20 siRNA knockdowns induced a visible and reproducible disruption in Golgi morphology (Fig.1.A and Supplementary Fig.1). The remaining levels of targeted protein were assessed by antibody staining against the protein. For each treatment, cells with more than 80% reduction in antibody intensity from the median of control cells were defined as having an efficient knockdown (analysis pipeline in Supplementary file 2). Our analysis revealed that a subset of cells with strong depletion of target protein still displayed seemingly an intact Golgi morphology (Fig.1.B) suggesting that transfection efficiency alone does not explain this inherent variability in morphological responses to siRNA mediated knock-down of the target genes. This motivated us to question the real reason for difference between the phenotypes, and to ascertain the mechanisms by which a subgroup of cells could prevent Golgi disruption. To this end, we developed a pipeline, shown in Fig.1.C, to correlate cells with different Golgi phenotypes under a single perturbation seen and marked by adaptive feedback microscopy, with their gene expression profiles at the single-cell resolution for a comprehensive comparison of the differences that underly distinct Golgi morphologies.

**Figure 1:**
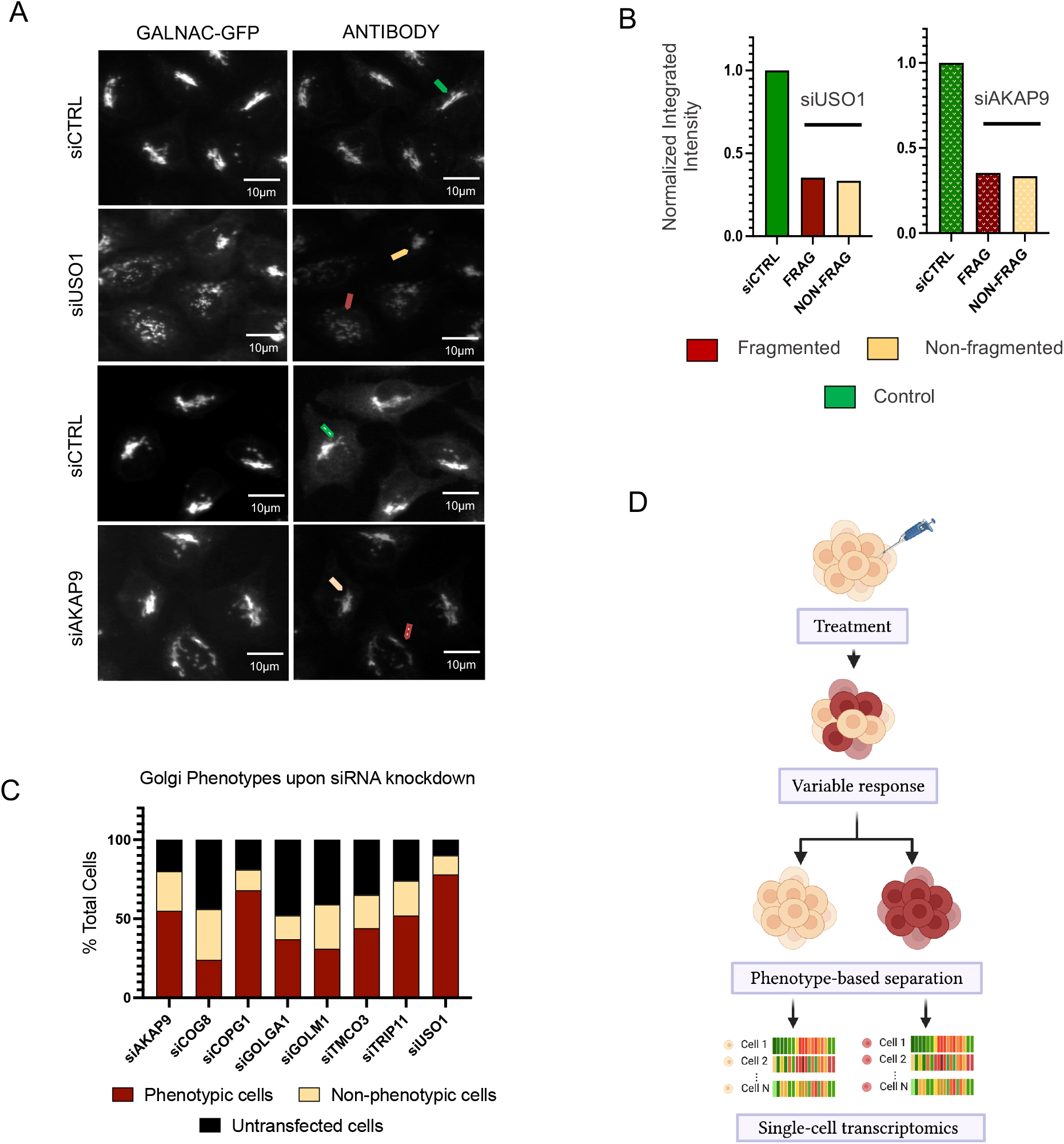
Differences in Golgi phenotypes upon siRNA mediated depletion. A) siRNA treated HeLa-GalNaC cells show a mixed responses in their Golgi morphology. Most cells display Golgi morphology characteristic of the treatment (red arrow), and a small proportion of cells with unperturbed Golgi morphology (beige arrow) B) Quantification of remaining protein levels by antibody staining shows similar levels of protein in cells with different Golgi morphologies (reg and beige arrows) as compared to cells transfected with control siRNA (in green). C) Quantification of Golgi phenotypes describing the percentage of cells which show siRNA knockdown displayed the phenotype typical of the treatment (red) as well as knocked down cells not showing the characteristic phenotype (beige). The black bar represents un-transfected cells (quantified by antibody staining) D) Description of a method to isolate single cells of a particular phenotype from a heterogenous population of cells to perform phenotype specific single cell transcriptomics.

To establish this pipeline, we chose USO1 also known as p115 as a candidate gene for siRNA knockdown and subsequent phenotype analysis. USO1 is an essential transport factor for COPII mediated vesicular traffic from the ER to the Golgi, as well as inter-cisternal transport within the Golgi^65,66^. It is shown to be critical for maintenance and *de novo* assembly of the Golgi complex, required for tethering of the incoming vesicle to its target membrane^67^. Depletion of this protein results in strong fragmentation of the Golgi complex. We observed this fragmentation in about 80% of USO1 depleted cells, whereas the remaining 20% displayed intact Golgi morphology despite USO1 depletion. These two observed populations are henceforth referred to as ‘fragmented’ and ‘non-fragmented’ phenotypes.

### Integrated adaptive feedback microscopy-based single-cell analysis pipeline for Golgi morphology

To be able to isolate these two populations after identifying them, HeLa-GalNaC-T2 cells were electroporated with a photo-activatable marker H2B-PAmCherry^68^ prior to a 72-hour siRNA mediated USO1 depletion (Supplementary Fig.2.A). This histone-tagged marker allowed us to selectively fluorescently label cells of interest by UV exposure to individual cells, converting fluorophores from dark to red. Golgi morphology of live cells after siRNA treatment was assayed by confocal microscopy experiments controlled by feedback analysis loop for identifying cells and selecting target phenotypes. Real-time image analysis was performed in either user-supervised or fully automated way to classify phenotypes as ‘fragmented’ or ‘non-fragmented’. The results of cell identification and phenotype selection were automatically communicated to the microscope acquisition software to generate individual ROIs on selected cells and immediately trigger their photo-activation (Supplementary Fig. 2.B). Cells of one phenotype were photo-activated per well of a culture plate, for each technical and biological replicate cells of both phenotypes were analysed. In addition, cells transfected with a control siRNA were also randomly marked using photo-activation. The wells were then trypsinized and sorted by fluorescent activated cell sorting (FACS) to collect single mCherry-positive (photo-activated) cells into a 96-well format (Supplementary Fig. 2.C). The collected cells were processed for single-cell RNA sequencing using the Smart-Seq2 protocol^69^. Resulting libraries were pooled after barcoding each single cell and sequenced at the Ilumina NextSeq500 platform.

### Transcriptome analysis shows USO1 depletion in both fragmented and non-fragmented cells

The single-cell transcriptome data was first subjected to quality control steps (Supplementary Fig 3.A) and sub-optimal batches were removed from further analysis. On average, 13,500 genes were detected in each batch (Supplementary Fig. 3.A). The resulting data was then analyzed for USO1 expression levels in both fragmented and non-fragmented single-cell populations. Transcriptomic data validated that both populations had USO1 significantly downregulated (Fig 2.A). The control population showed a wide range of USO1 expression values, presumably due to normal cycling of the protein’s levels with cell cycle. Next, the cell cycle phase of each cell was determined by using the sequencing data as described by Tirosh l. et al^70^ as well as by flow cytometry analysis. Both analyses showed similar results (Fig 2.B), which revealed a comparable proportion of cells in each of the cell cycle phases in both phenotypes. Both phenotypes showed a slight difference from the control population, which had a higher proportion of cells in S phase However, this difference was not statistically significant.

**Figure 2:**
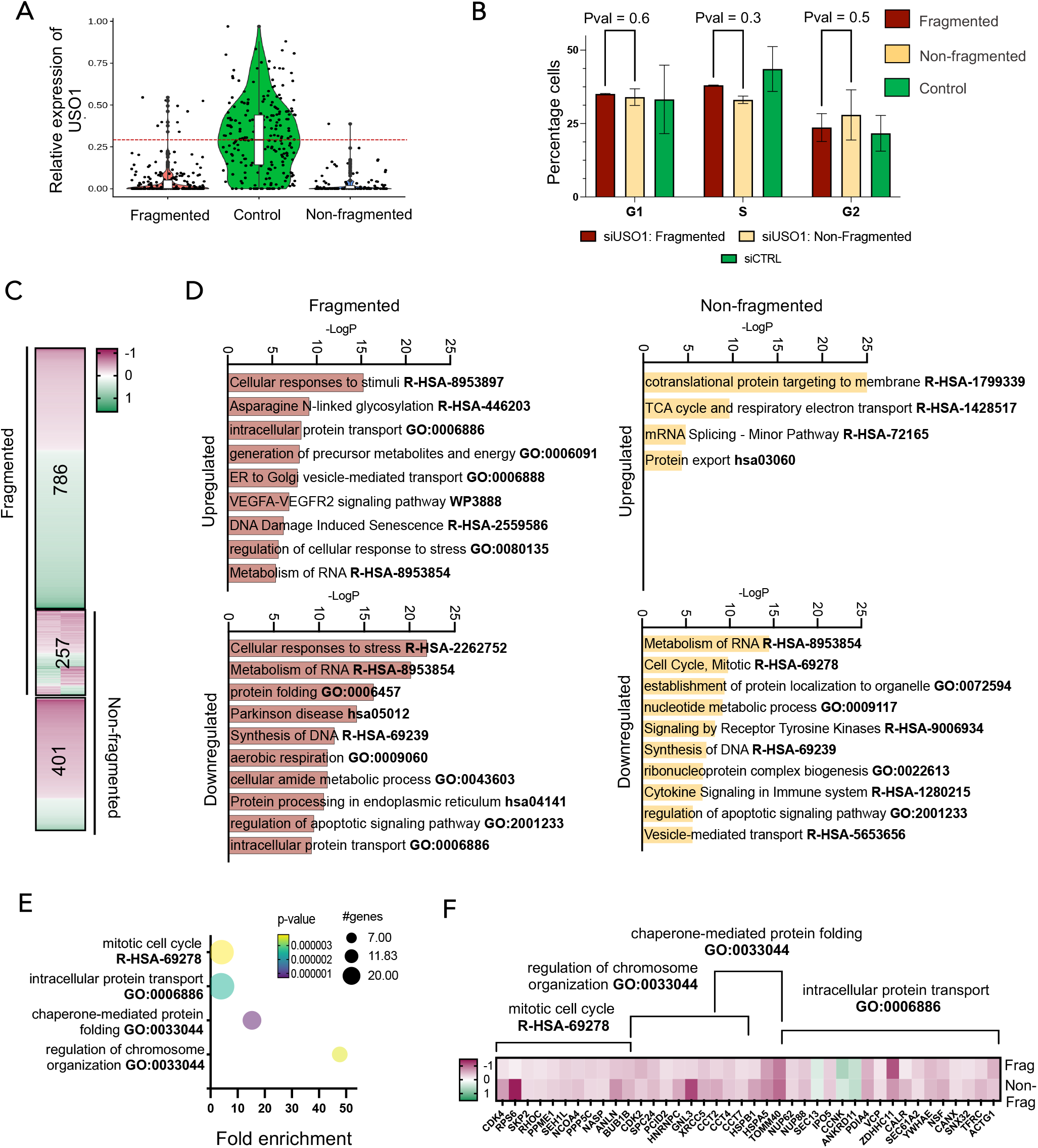
Single-cell transcriptome data describing the effect of USO1 knockdown. Relative USO1 expression in fragmented and non-fragmented phenotypic and control (control siRNA) shows a strong depletion of USO1 mRNA in both phenotypes, each dot representing a single cell. Cells treated with negative control siRNA show a wide range of expression values. B) Cell cycle analysis shows that both phenotypes have similar percentage of cells in each phase of the cell cycle, which did not differ significantly between phenotypes (Pval = 0.6 in G1, 0.3 in S and 0.5 for G2 cells). The differences to control cells were also not significant (not shown here). C) Heatmap summarizing differential expression analysis of both phenotypes’ v/s control cells. 1044 genes are DE in the fragmented phenotype and 658 genes in the non-fragmented phenotype. 257 of these are common to both phenotypes. D) Pathways represented by the genes differentially expressed in each phenotype v/s control cells are shown in respective bar graphs. E) Pathways represented by the 257 DE genes shared by both phenotype as common response to USO1 knockdown. F) Key genes in the pathways described in Fig. 2.E with the Log2 fold change in expression in each phenotype normalized to control cells are visualized in the heatmap.

Differential expression (DE) analysis of each phenotype versus control cells was performed to ascertain the effect of USO1 depletion on either phenotype (full list in Supplementary tables 3 & 4). 1044 genes were differentially expressed in fragmented cells, and 658 genes in non-fragmented cells (Fig.2.C). Functional enrichment analysis of the genes unique to each phenotype (up/downregulated) is shown in Fig.2.D. The fragmented phenotype showed an upregulation of genes involved in glycosylation, ER-Golgi transport as well as metabolism, among others. Genes involved in cellular stress response, and protein folding were downregulated. In the non-fragmented phenotype, genes pertaining to pathways of SRP-mediated co-translation, TCA cycle and protein export were upregulated, whereas genes involved RNA metabolism, mitosis and protein localization were downregulated. 257 genes were common to both phenotypes, and Fig 2.E shows the main pathways represented by these genes. Four main processes show enrichment in response shared between phenotypes (complete analysis is in Supplementary tables 5-8). These include cell cycle, intracellular protein transport – which have a higher number of genes in our dataset, but a lower fold enrichment. Chaperone mediated protein folding, and chromosome maintenance genes show significant fold enrichment, albeit with a few key genes present in the shared response by both phenotypes. The specific genes found are shown in Fig. 2.F along with their Log2 fold change in comparison to control cells. Worth noting, all the processes found were seen to be downregulated in USO1 depleted cells in comparison to control cells.

### Fragmented and non-fragmented phenotypes differ in unexpected trafficking pathways

Having ascertained a transcriptome-wide response to the treatment, we now performed DE analysis between the two phenotypes. In total, 550 genes were found differentially expressed between the two phenotypes (adjPvalue < 0.1), of which 461 were up regulated in fragmented cells compared to non-fragmented cells (Fig. 3.A, full list in Supplementary table 9). Clustering of differentially expressed genes between the two phenotypes into biologically relevant pathways revealed an upregulation in fragmented cells predominantly of pathways pertaining to cellular localization, protein transport and regulation of apoptosis (Fig. 3.B). A closer look at the genes conceded a range of different pathways, the most striking of which were post-Golgi transport to the endosomal/lysosomal network and plasma membrane (Fig. 3.C). Also of interest were several genes pertaining to the regulation of mRNA export and splicing. Surprisingly, very few genes of the targeted pathway, i.e., ER to Golgi transport, had significantly different expression between the two phenotypes. As shown in Fig 3.D, barring a handful, all DE pathways seemed consistently downregulated in the non-fragmented phenotype, with control levels or slight upregulation in fragmented cells (complete analysis in Supplementary table 10).

**Figure 3:**
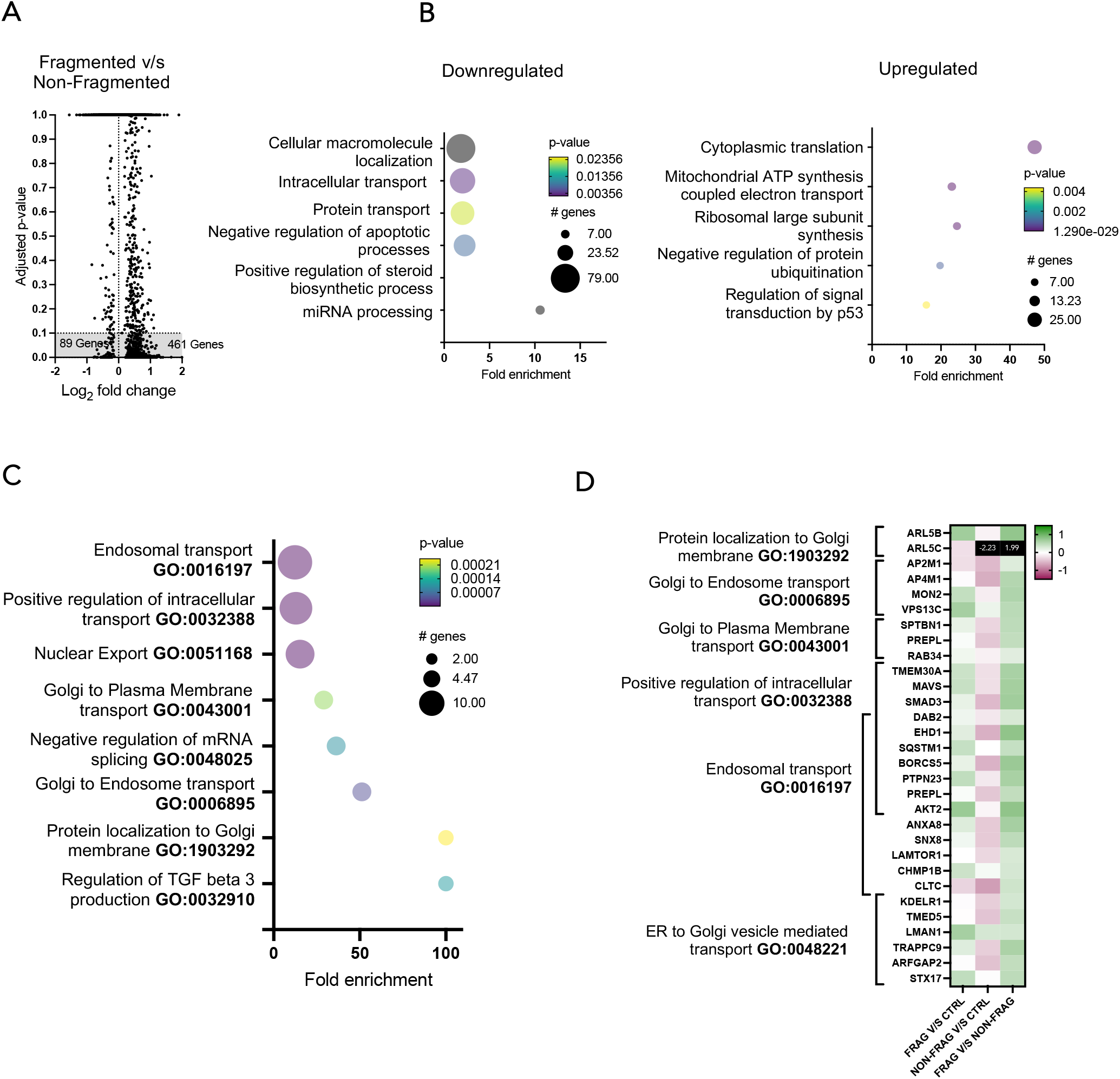
Differentially expressed genes in Fragmented v/s Non-fragmented phenotypes. A) Volcano plot showing differential expression between Fragmented v/s Non-fragmented phenotypes. 550 genes (grey shaded area) are significant and taken for further pathway analysis. B) Pathway enrichment analysis of genes downregulated (left) and upregulated (right). GO term of the enriched pathway is on Y-axis, and X-axis shows fold enrichment of the pathway in the DE gene set. The size of the bubble represents the number of genes and color of the bubble indicates the p-value of the enrichment. C) Detailed classification of the GO term ‘protein transport in Fig. B shows enriched pathways relating to post-TGN and endosomal transport. GO term of the enriched pathway is on Y-axis, and X-axis shows fold enrichment of the pathway in the DE gene set. The size of the bubble represents the number of genes and color of the bubble indicates the p-value of the enrichment D) Key genes in the pathways described in Fig. 3.C with the Log2 fold change in expression in each phenotype normalized to control cells, as well as the DE expression values between fragmented and non-fragmented phenotypes.

Most of all, the finding of post-TGN trafficking pathways being downregulated in non-fragmented cells involving key components - such as clathrin heavy chain (CLTC), adaptor proteins AP2M1 and AP4M1 was very unexpected. This led us to consider the possibility of the phenotypic differences arising from changes in the late secretory and endocytic pathways. If these genes played a decisive role for the phenotype, their knockdown in combination with USO1 might rescue USO1 induced fragmentation. To examine this, co-knockdowns of USO1 were performed with the strongest effectors in our gene list-AP2M1, CLTC and ARL5C. Remarkably, AP2M1 and CLTC co-knockdowns with USO1 compensated the fragmented phenotype caused by USO1 knockdown alone (Fig.4.A). We did not observe a significant difference with ARL5C co-knockdown, where cells display a predominantly fragmented Golgi. For comparison, we performed a co-knockdown of a gene operating in ER to Golgi transport, namely TMED5. This resulted in re-distribution of GalNaC signal back into the ER accompanied by fragmentation in many cells (Fig.4.A). Singular knockdowns of AP2M1, ARL5C and TMED5 did not give rise to a perturbed Golgi phenotype visible by light microscopy, whereas CLTC downregulation alone resulted in mild Golgi fragmentation. This significant rescue of phenotype signals that the endocytic proteins may indeed play a vital role in maintenance of Golgi architecture. Fig. 4.B outlines the pivotal pathways and key players that show decreased expression in non-fragmented cells, the possible implications are discussed below.

**Figure 4:**
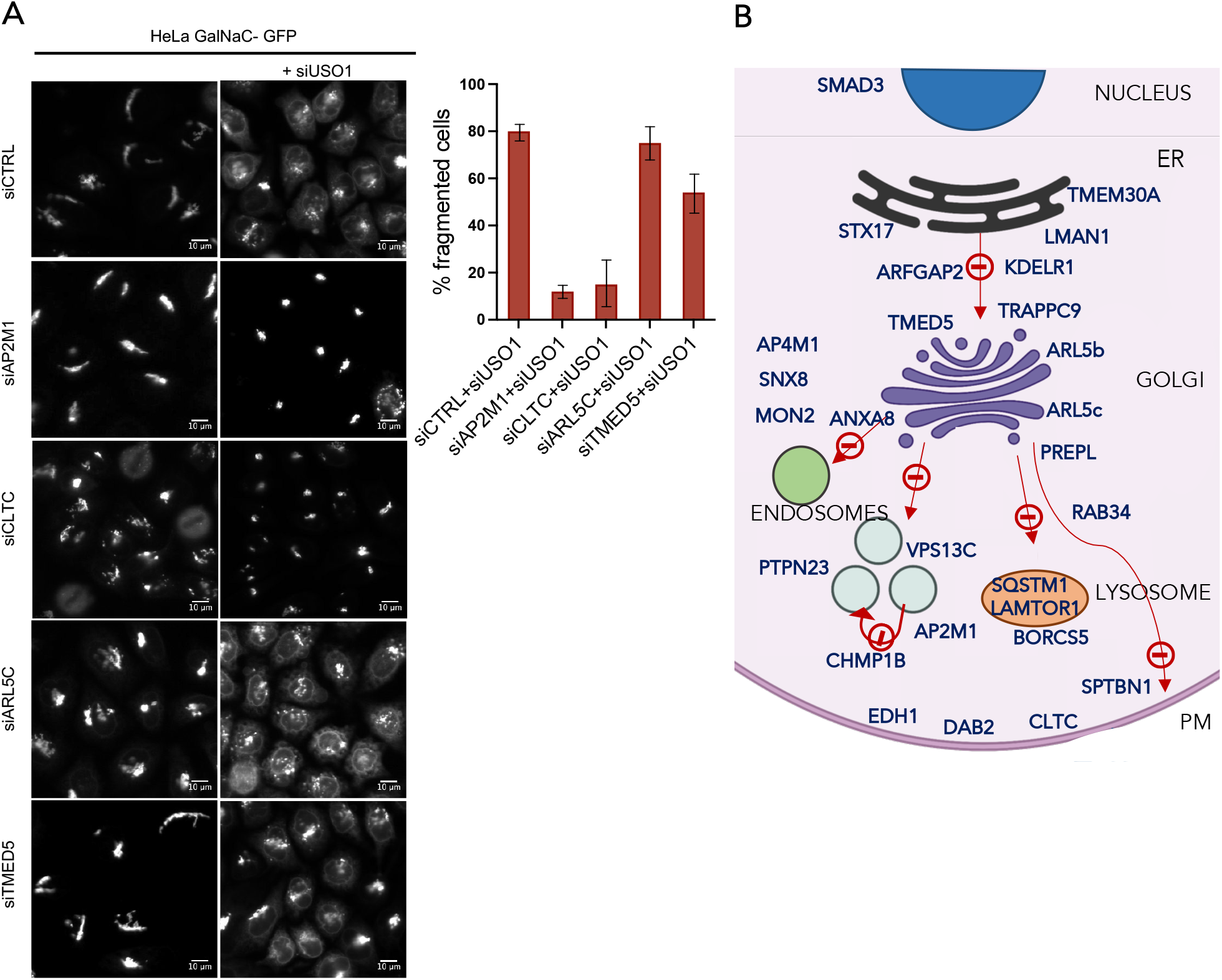
Endosomal and endocytic pathways may underline phenotypic differences. a) Co-knockdowns of USO1 with four hits in the DE analysis between phenotypes show rescue of the fragmented phenotype by AP2M1 and CLTC but not ARL5C and TMED5. Left panel shows individual siRNA knockdowns of the four genes in HeLa-GalNaC-T2 cells with the Golgi complex visualized. Right panel shows the changes in Golgi morphology when the four genes are knocked down respectively in combination with USO1 knockdown, also siRNA mediated. The bar graph on the right shows the percentage of cells with fragmented Golgi phenotype resulting from each of the co-knockdowns. B) Schematic of the secretory pathway outlining the genes identified in our dataset as significantly downregulated in non-fragmented cells as compared with fragmented cells.

## Discussion

The structural maintenance of the Golgi complex has been a long-standing question in cell biology, a complete understanding of which is made imperative by the mounting evidence of the organelle as a regulatory platform for multiple signaling pathways. We employed a unique approach to this problem, taking advantage of intrinsic variability in Golgi structural responses to siRNA depletion. To this end, we developed an adaptive feedback microscopy pipeline that integrates high quality imaging, fast targeted photoactivation and online image analysis for high-throughput phenotype identification and marking for subsequent cell sorting. This approach has the advantage of higher spatial resolution, required for correct classification of Golgi phenotypes, over imaging flow sorters^71^ – an alternative strategy for such multimodal analysis, where image quality is often compromised. Other successful approaches have also been described in literature, albeit requiring specialized equipment/code for cell collection^72,73^. Marking of selected cells using photoactivation followed by flow cytometry is a simplistic way of collecting cells without the use of complex microfluidics chips and precision tools described in other approaches with the same goal^74,75^, thereby making this pipeline accessible to many labs. A possible caveat for expanding our pipeline is the limited number of colors (fluorophores) that can be used to isolate multiple phenotypes in the same sample. Although we used this pipeline to obtain transcriptional responses of our phenotypes, this pipeline remains compatible with practically any other approach that provides a single-cell or low-input readout (genomics, proteomics, etc.). Our pipeline to correlate phenotypic response with single-cell transcriptomics is the first method of its kind used to study Golgi organization and has led to several key findings that further our understanding of the organelle regulation as follows.

First and foremost, we show at the single-cell USO1 mRNA quantity reduced to similar levels for both phenotypes, which was in coherence with our observations at the protein level determined by antibody staining. This validates our hypothesis that the difference in phenotypes is not merely an effect of poor transfection efficiency. Another common explanation of phenotypic variability could be difference in cell cycle phases, where we also showed using both transcriptome data and flow cytometry analysis that both phenotypes show very similar percentages of cells in each cell cycle phase. It is worth noting that both phenotypes differed in cell cycle marker expression from control cells, although not significantly. Cell cycle genes also showed up strongly as a shared response between phenotypes, supporting the analysis from earlier. This is consistent with the literature, as USO1 has been shown to be important for cell division, in particular proper chromosome segregation and cytokinesis^76^. It is also regulated in a cell cycle dependent fashion by phosphorylation, wherein the phosphorylated form is found predominantly in interphase cells and the dephosphorylated form is observed in dividing cells.^67^

### Individual response of each phenotype reveals expression differences of ER-Golgi trafficking genes

Transcriptome data revealed about 800 genes differentially expressed exclusively in the fragmented phenotype. The non-fragmented phenotype had less than half that number, with about 400 genes unique to the phenotype. Strikingly, cells of the fragmented phenotype showed upregulation of several transport genes, such as SEC24A- a COPII component, COPG1 -a COPI component as well as SEC22B, KDELR2/3, YIPF4, YIF1B, all of which are involved in ER to Golgi anterograde or retrograde transport. It has been shown in the literature that depletion of USO1 inhibits transport of some cargo, particularly the Vesicular Stomatitis Virus Glycoprotein (VSV-G)^77^, but has limited effects on soluble cargo in many cell lines.^78^ The increased expression of these genes in fragmented cells can be seen as an attempt to restore any ER-Golgi transport defects/delays caused by USO1 depletion. The same genes are however expressed at control levels in the non-fragmented phenotype. This implies that a different mechanism is being used by non-fragmented cells to compensate for possible trafficking defects, which means that the morphology of the Golgi has a role of either the cause or consequence of this compensation mechanism.

### Shared response between phenotypes reveals Cell Cycle genes downregulated in USO1 depleted cells

Aside from the unique responses to USO1 depletion, both phenotypes shared 257 genes which had a common response to USO1 depletion. We saw downregulation of several genes involved in cell cycle progression and mitosis shared between phenotypes versus control cells. These included those essential for G1/S phase transition, such as CDK2 and CDK4, and those functioning in chromosome segregation, such as SPC24, SEH1L, ANLN, BUB1B and NUP62. Additionally, we observed genes required for proper protein folding as part of the chaperonin-containing T-complex (TRiC), with an enrichment of protein folding required for telomere maintenance. The gene encoding for Valosin Containing Protein (VCP), necessary for the fragmentation of Golgi stacks before mitosis, was surprisingly also downregulated in both phenotypes. These findings suggest that the functions of USO1 in the context of cell division might be independent of Golgi morphology presentation. The fact that the non-fragmented phenotype displays intact morphology, does not mean a functional rescue of the knockdown. The knockdown of USO1 also had a minimal effect on secretory pathway proteins, with a few ER to Golgi transport proteins showing downregulation, such as NSF, SNX32 and SEC61A2. In addition, there was a complete absence of genes relating to post Golgi trafficking. We anticipated to find direct interactors of USO1, of which prominent ones are GM130, Rab1, GMAP210 and other SNAREs^79^ as well as members of the COG ^80^ complex, all of which thought to be important in maintenance of the Golgi complex via their interaction with USO1. Much to our surprise, this was not the case. No direct interactors of USO1 were found to change expression between the phenotypes or between either phenotype versus control cells, suggesting that their individual functions are not dependent on the presence of USO1.

### Differential response of the two phenotypes reveals differences in post-TGN trafficking genes

The differential expression (DE) analysis between phenotypes revealed 550 genes of significantly different expression, corresponding roughly to 4% of all detected genes. This is a striking finding, given that the cells were distinguished solely based on Golgi morphology with all other parameters (culture conditions/ siRNA treatment) being held constant. This not only demonstrates the much wider implication of a single gene silencing experiment on the cell, it also shows that important biological information might be gained from exploiting phenotypic heterogeneity.

DE analysis between the two Golgi phenotypes showed predominant upregulation of pathways pertaining to cellular transport, regulation of apoptosis and miRNA processing in fragmented cells versus non-fragmented, which turned out to be in fact downregulation of these pathways in the non-fragmented cells. Much to our surprise, several genes operating in post-Golgi transport pathways were strongly downregulated in non-fragmented cells. This included important genes in endosomal transport such as clathrin heavy chain (CLTC), clathrin adaptor proteins (AP2M1, AP4M1), and many others. The unanticipated discovery of late secretory pathway genes differentially expressed between phenotypes implies that phenotypic differences are linked to trafficking related functions of USO1. Moreover, it points towards the presence of a feedback sensing mechanism that whereby the late and early secretory pathways communicate upon an error in normal trafficking. Rescue of Golgi fragmentation upon co-knockdown with AP2M1 and CLTC but not TMED5 further strengthens this hypothesis. Non-fragmented cells may rely on to a dampening of post-Golgi traffic and endocytosis to maintain homeostatic pools of essential material such as membrane receptors at each organelle in the absence of fresh cargo. It is difficult to say if the two phenotypes arise stochastically at different rates, the fragmented one dominating in the case of USO1, or if there are underlying cellular states that predetermine the compensation mechanism adopted. Although the traditional view is to think of Golgi morphology being a result of a perturbation, changes in Golgi morphology may also considered as a way to regulate cellular responses. Although an exhaustive analysis of all genes/pathways found in our DE analyses (Supplementary tables 3-8) is beyond the scope of this paper, our pipeline illustrates how a relatively simple approach can add more value to cellular transcriptomic data by the added the dimension of morphological information.

## Methods

### Cell culture, electroporation and siRNA transfection

HeLa GalNac-T2 cells were cultured in low glucose (1g/l) DMEM (Gibco) supplemented with 10% (w/v) foetal calf serum and 2mM L-Glutamine. 400μg/ml Geneticin was added to the cells to maintain selection pressure of the GFP tagged enzyme and cells were kept at 37°C and 5% CO2. The photo-activatable plasmid H2B-PAmCherry was transiently expressed in HeLa GalNaC-T2 cells in suspension by electroporation. Electroporation mix was performed using the Lonza Amaxa SE Nucleofection 4D kit in the 100μl format with the 4D-NucleofectorTM X Unit (Lonza) machine. For subsequent siRNA transfection, electroporated cells were plated one day prior to transfection on 24-well glass bottom plates at a density of 15,000 cells per well. siRNA transfections were performed using pre-designed siRNAs against the target transcripts (full table in supplementary) at a working concentration of 30μM. Oligofectamine™ was used as the transfection reagent, and both siRNA and Oligofectamine were diluted in OptiMEM before mixing them together. Prepared mixes were added to cells under starved conditions (FCS free media) and incubated for 4 hours at 37°C, followed by addition of media containing 2X FCS. The cells were taken for further analysis/imaging 72 hours after transfection. Imaging of fixed cells was performed at the Olympus ScanR with the 20x UPlanApo NA 0.7 Air Objective. Three fluorophores were imaged-DAPI (Nuclei), GFP (Golgi) and A568 (antibody staining), with 50 fields of view taken per treatment condition.

### Image analysis

The offline image analysis for quantifying fractions of different phenotypes and measuring protein levels on the images of fixed cells was done using a Cell Profiler pipeline that can be viewed in Supplementary table 2.

### Live-cell feedback microscopy and photoactivation

Imaging of live cells and photo-activation of selected phenotypes was performed on the Zeiss LSM900 Confocal microscope controlled by the ZenBlue software. The acquisitions were performed with either Plan-Apochromat 20x/0.8 M27 air objective (FWD=0.55mm) or Plan-Apochromat 40x/0.95 Corr M27 air objective (FWD=0.25mm). The photoactivation of the target regions was performed with confocal scanner using 405 nm laser. In our experiments 30-40 scanning iterations per region with 2% of 405 nm laser and pixel dwell time of 0.61 μs were required to achieve the optimal photoactivation efficiency. Although one might expect to perform faster photoactivation by using higher laser powers and faster scanning speeds, which are technically possible, in practice moderate laser powers need to be used to avoid bleaching of photoactivated fluorophores.

To facilitate the photoactivation of many cells in multiple positions in the well plates, we set up the adaptive feedback microscopy pipeline. Three imaging jobs were acquired at each stage position. First, low resolution stack containing 51 z-planes was acquired the Hoechst channel using 405 nm laser excitation at low power (Supplementary Fig.2B.1). This stack was then automatically processed by online image analysis to define the best focus plane for the cells in the field of view (Supplementary Fig.2B.2). Second, high resolution imaging of the field of view (319.45 × 319.44 μm with the pixel size of 0.2507 × 0.2507 μm) was automatically triggered in Hoechst, GFP and mCherry channels. This image was then subjected for image analysis for segmenting cells and selecting target phenotypes (Supplementary Fig.2B.3,4). Two options for the image analysis were developed. The fully automated analysis identifies the cells and selects those of the right phenotype without any user supervision. The semi-automated image analysis enquires user to mark the positions of the target cells. For single cell transcriptomics experiments we used the semi-automated routine, fully automated pipeline was also tested and validated. For both pipeline versions the automated transfer of identified positions to the microscope triggered the third image acquisition job, which included photoactivation of the target regions and acquiring post-activation images in three channels for documenting photoactivation results (Supplementary Fig.2B.5).

The image acquisition and the photoactivation in the adaptive feedback microscopy pipeline was run by the dedicated macro in ZenBlue software, and online image analysis was established using AutoMicTools library for Fiji previously developed by us. The developed AutoMicTools pipeline, ZenBlue macro and the instructions for running the experiments are publicly available (https://git.embl.de/grp-almf/automictools-photoactivation). The developed pipeline can be easily adopted to automatically photoactivate different phenotypes. For this purpose we included the possibility to incorporate into the pipeline custom Fiji script implementing segmentation and phenotype identification for particular project.

### FACS sorting and single-cell sequencing

After photo-activation of the two different phenotypes in different wells of a glass-bottom 24 well plate (Celvis), cells were detached from the glass bottom well using TrypLE™ Express (ThermoFisher) and resuspended in complemented DMEM to make up the volume of the suspension to 1ml in an Eppendorf tube. The cell suspension was centrifuged (5417R) at 1000 rpm at 4°C for 5 minutes to obtain a cell pellet. The cell pellet was washed with cold FACS wash buffer and centrifuged as before. The supernatant was discarded, and the pellet was resuspended in 250μl cold FACS staining buffer (PBS, 0.2% BSA, 0.1% saponin,1:100 RNasin). Samples were kept on ice to prevent cell lysis. The cells were sorted using the BD FACS Aria as single cells into a 96-well plate (Bio-Rad) containing 4μl lysis buffer (0.4% Triton X-100 in PBS, 1:20 (40U/μl) RNasinPlus, 10mM dNTP mix, 5 M Oligo dT) in each well. The gating on the sorter was set up to first select for live cells using Hoechst staining and then to select for GFP and mCherry double positive cells determined by negative controls for GFP and mCherry respectively. Thus, only photo-activated cells expressing the GFP-tagged Golgi marker used for phenotype identification were collected. About one 96-well plate (96 cells) were collected per phenotype per experiment.

Libraries were prepared with SMART-Seq protocol and sequenced on NextSeq 500 instrument.

### Differential expression analysis

Single cell reads were mapped with STAR (version 1.15.1) on GRCh38 genome. Gene count tables were also computed during the alignment step. Cells expressing a gene count less than 4500 and a percentage of mitocondrial gene expression higher than 25% were filtered out. The read counts of the 705 selected cells (273, 264 and 168 for fragmented, Neg8 control and non-fragmented respectively) were imported into Seurat R package (version Seurat_3.1.5). Cells were analyzed and differential expression analysis of both populations against the control was done with the same package. Functional enrichment analysis was done using Cytoscape with the add-ins Cluego (with GO term fusion), as well as with Metascape and Panther web tools for FDR calculation and Bonferroni correction for multiple testing.

