## Supplementary Figures for "Phenotype-based single-cell transcriptomics reveal compensatory pathways involved in Golgi organization and associated transport"

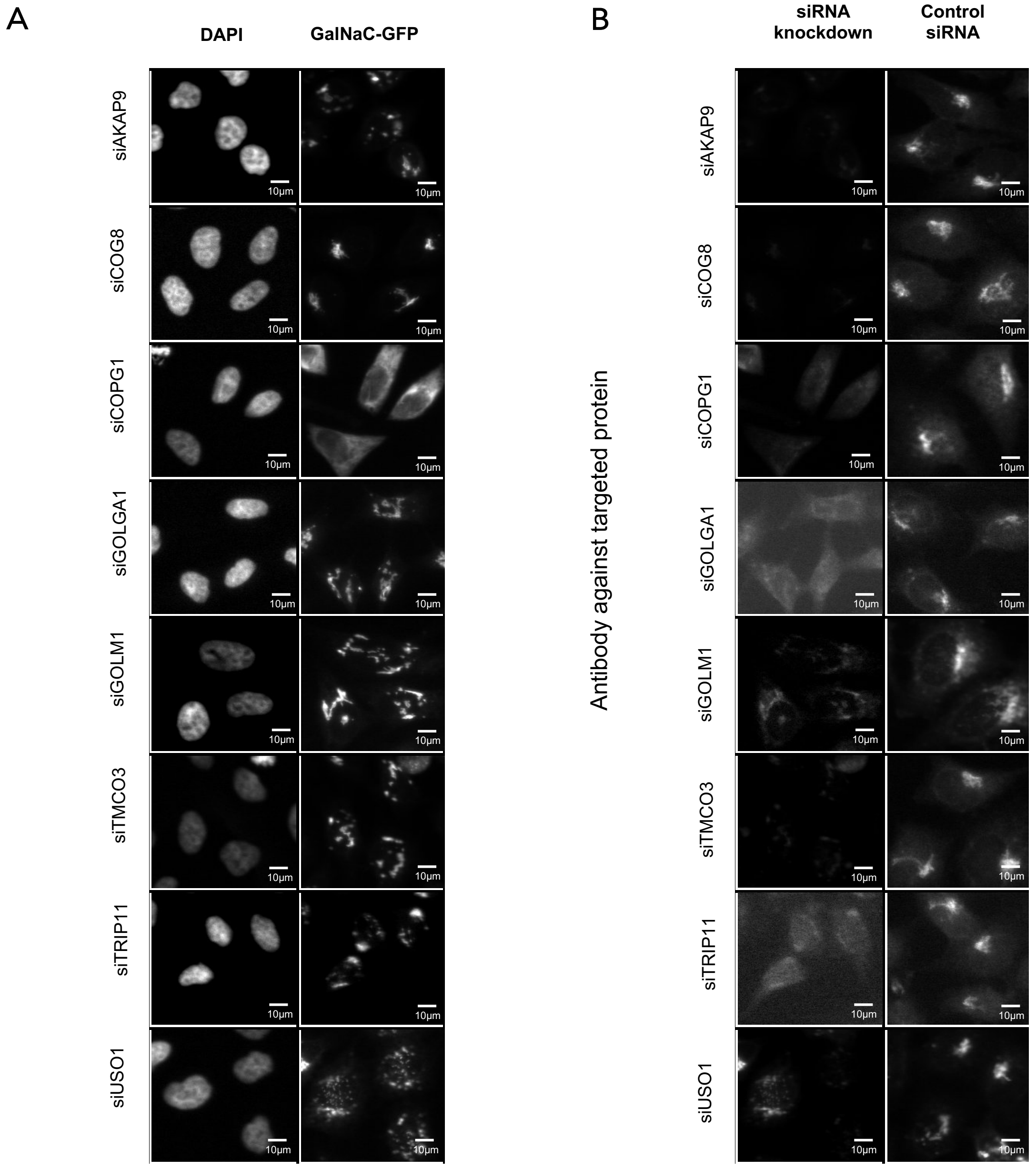


<u>Supp. Fig. 1 siRNA knockdown and resulting Golgi phenotypes of 8 proteins.</u> A) Golgi phenotypes upon knockdown of several Golgi localized proteins. Shown here in DAPI are the nuclei, and the Golgi complex is seen by the GalNaC-GFP signal in the right column. B) Remaining levels of the targeted mRNA are observed using antibodies specific to the target protein and compared side by side with protein levels in control siRNA treated cells.

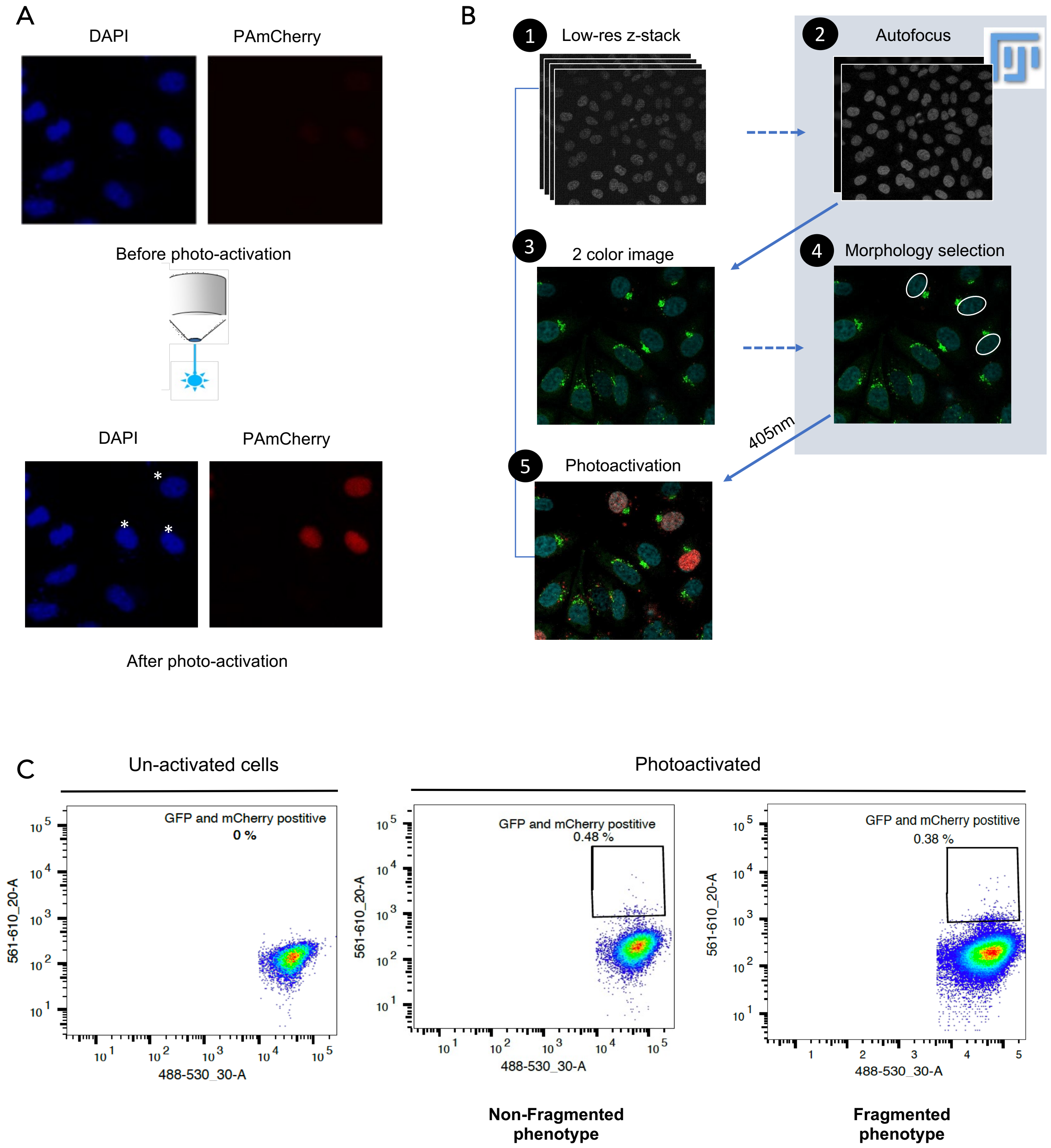


Supp. Fig 2: Pipeline for phenotype-based single-cell transcriptomics. A) Photoactivation using H2B tagged PAmCherry fluorophore that is electroporated/transfected into cells. UV exposure of nuclei at 405nm results in dark (inactive) to red (active) conversion of the fluorophore. B) Golgi phenotypes are identified using online image analysis in Fiji. The feedback microscopy pipeline triggers this image analysis and uses its results to photoactivate the selected phenotypes. B.1 shows a low-resolution z-stack, followed by an autofocus performed in Image J (2). Images 3, 4 and 5 show part of FOV of the high-resolution image showing the Golgi, phenotype selection and subsequent photoactivation, respectively. C) Photo-activated cells are then collected as single-cells for RNA-seq using FACS. The left graph shows the flow plot of an un-activated sample, used for gating. The center and right graphs show the gates used to collect activated cells in the two phenotypes. The log intensity of the GalNaC-GFP (488nm) signal is plotted on the X-axis and the Y axis denotes the log intensity of the activated PAmCherry fluorophore (561nm).

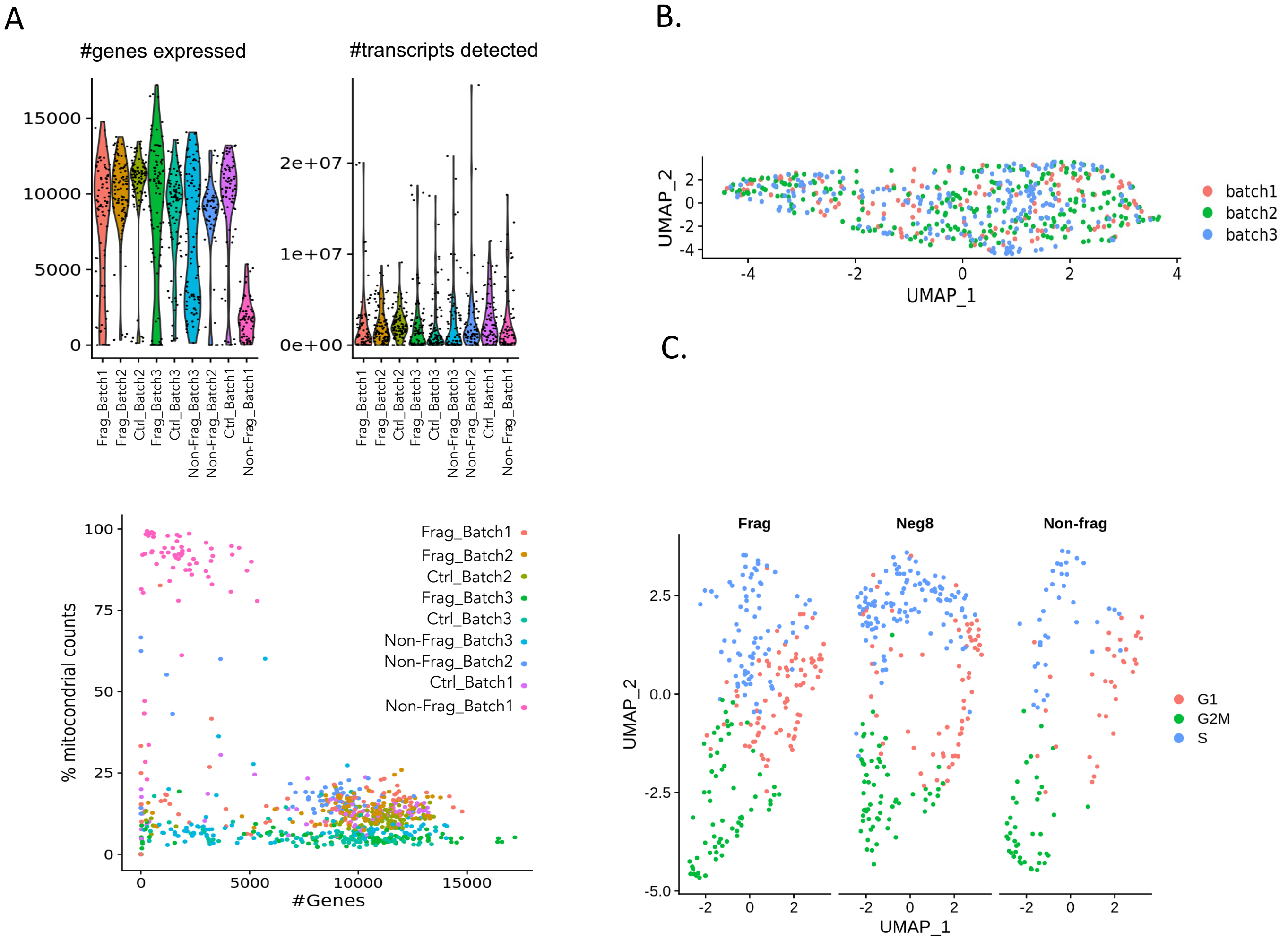


Supp. Fig 3: Quality control of single-cell transcriptomic data. A) Plots representing the number of genes and transcripts detected in each experimental batch. Each dot represents a single cell. X-axis represents the experimental batch and phenotype. The lower graph shows the percentage of mitochondrial counts with color representing the experimental batch. Non-fragmented cells of batch 1 were excluded from further analysis due to high mitochondrial counts, suggesting damaged cells. B) UMAP graph of the three batches show a homogenous distribution ruling out batch effects. C) UMAP graph depicting the number of cells in G1, G2M and S phase of the cell cycle in red, green and blue, respectively. Each dot represents a single cell.
